# VarSAn: Associating pathways with a set of genomic variants using network analysis

**DOI:** 10.1101/2020.12.22.424077

**Authors:** Xiaoman Xie, Matthew C. Kendzior, Xiyu Ge, Liudmila S. Mainzer, Saurabh Sinha

## Abstract

There is a pressing need today to mechanistically interpret sets of genomic variants associated with diseases. Here we present a tool called ‘VarSAn’ that uses a network analysis algorithm to identify pathways relevant to a given set of variants. VarSAn analyzes a configurable network whose nodes represent variants, genes and pathways, using a Random Walk with Restarts algorithm to rank pathways for relevance to the given variants, and reports p-values for pathway relevance. It treats non-coding and coding variants differently, properly accounts for the number of pathways impacted by each variant and identifies relevant pathways even if many variants do not directly impact genes of the pathway. We use VarSAn to identify pathways relevant to variants related to cancer and several other diseases, as well as drug response variation. We find VarSAn’s pathway ranking to be complementary to the standard approach of enrichment tests on genes related to the query set. We adopt a novel benchmarking strategy to quantify its advantage over this baseline approach. Finally, we use VarSAn to discover key pathways, including the VEGFA-VEGFR2 pathway, related to de novo variants in patients of Hypoplastic Left Heart Syndrome, a rare and severe congenital heart defect.

## INTRODUCTION

The relationship between genotypic differences, e.g., single-nucleotide polymorphisms (SNPs), and health-related differences among individuals is a major topic of research today. A common approach is to find polymorphisms (variants) that are statistically correlated with phenotypic differences, as in genome-wide association studies (GWAS) (1) and family-based association tests (2). An alternative approach involves “trio-based” genome sequencing (3,4) of modest-sized cohorts of patients to identify de novo mutations in the child, and genes and pathways that are frequently impacted by these variants. Irrespective of the specific methodology used, it is common for genotype-phenotype studies to identify a set of SNPs, promising to reveal mechanistic insights into phenotypic variation. Commonly, individual SNPs or genes related to them are subjected to further experimental interrogation (5),and there is intense on-going research into efficient ways to shortlist variants for experimental follow-up (6–10). In parallel, researchers often seek to understand the biological processes that are implicated by the identified SNPs (11). Our work addresses this latter goal. In particular, we present a novel computational tool to answer the following bioinformatics question arising from genotype-phenotype studies: *given a set of SNP positions of potential relevance to a phenotype, which pathways are likely to be important to the phenotype?* We refer to this as the “variant set characterization problem”.

The variant set characterization problem is conceptually similar to the highly popular “gene set characterization” problem, where a given gene set, e.g., differentially expressed genes from a transcriptomic study, is tested for statistical association with biological pathways, Gene Ontology terms, etc. (12). Indeed, one solution to the variant set characterization problem is to map SNPs to genes, e.g., based on predicted impact of a SNP on the function of a gene’s encoded protein, and then subject the resulting genes to the plethora of available gene set characterization methods (11), including enrichment tests (13,14) and pathway analysis tools (15). Of particular relevance to our work is the class of methods that test for associations between the study-based gene set (in this case, derived from the phenotype-related SNP set) and pre-determined gene sets from a compendium such as REACTOME (16), KEGG (17) or Gene Ontology (18). Commonly, this is done via a Hypergeometric test of overlap between the two gene sets, although alternatives that make use of rankings (rather than merely sets) of genes have also been widely used (19,20).

Here, we focus on the task of associating a given SNP set with pre-determined gene sets. Specifically, the given SNPs are not assumed to be ranked or assigned numeric scores, and the candidate pathways with which the SNPs may be associated are not assumed to have a known network structure or “topology” (12). These decisions were motivated by considerations of simplicity and wider applicability. Admittedly, in many cases these assumptions may result in loss of valuable information, e.g., GWAS SNPs have associated p-values that may be used to score or rank them in the course of finding the most relevant pathways (21), and pathway topology may be used to good effect in gene set characterization approaches (22–25). Nevertheless, we tackle here the simpler version of the variant set characterization problem because it allows the proposed method to be used even when SNP scores are not available (e.g., de novo variants from a trio-based studies) and gene sets defined by a common annotation (e.g., a Gene Ontology term or the targets of a transcription factor) are used in place of pathways.

As noted above, the most natural solution for the particular problem posed above is via a test of enrichment between the SNP-related genes and pathway genes: if many of the given SNPs point to genes that belong to the same pathway, the pathway is considered relevant to the given SNP set. The intuitive appeal and statistical rigor of this approach are reasons in its favor. Yet, some aspects of the approach merit closer inspection and potential changes: Firstly, the step of mapping the given SNPs to genes raises questions. For instance, a SNP may be mapped to a gene if it is predicted to impact the protein function (26), or if it is a cis-regulatory eQTL (expression quantitative trait loci) of the gene (6); in either case the gene’s membership in a pathway furnishes evidence for the relevance of that pathway. But if both types of SNPs (coding and non-coding) are present in the given SNP set, do their mapped genes provide equal evidence for relevance of respective pathways that include those genes? Even among non-coding SNPs, should a gene related to a SNP based on eQTL evidence be considered as valuable as a gene to which a SNP has been mapped purely based on proximity? Furthermore, if one SNP points to one gene and (say) three other SNPs point to another gene, should the two mapped genes have equal importance in determining relevant pathways, or should the latter gene receive thrice as much importance?

A second class of questions arises in the gene-to-pathway mapping step. The enrichment-based approach counts the number of mapped genes that belong to a pathway. But if a mapped gene belongs to, say, 10 different pathways, and another mapped gene belongs to a unique pathway, should we consider the latter pathway as more relevant to the gene set (and hence to the SNP set) as there is more specific evidence in its support? Moreover, if a mapped gene is not annotated as a member of a pathway, but is known to interact with another gene (e.g., by physical interactions between the encoded proteins) and the latter gene belongs to a pathway, should this be considered as (at least partial) evidence of the relevance of the pathway? There is some work showing (27) that appropriate consideration of such indirect evidence of a gene’s relationship to a pathway is beneficial to pathway analysis.

The above considerations motivated us to develop a new computational method to solve the variant set characterization problem. The new tool, called VarSAn (<u>Var</u>iant <u>S</u>et <u>An</u>notator, pronounced “version”), builds on the intuitive appeal of the enrichment-based approach of counting SNP-related genes that belong to the same pathway, but re-casts the problem in a network-analysis framework that addresses the above-mentioned concerns. We employ VarSAn to analyze sets of GWAS SNPs and somatic mutations related to breast cancer (BrCa) and prostate cancer (PCa), as well as GWAS SNPs related to drug response variation. Through these applications we demonstrate the effects and advantages of key methodological choices in the tool, e.g., identifying relevant pathways that are supported by the literature but are not ranked highly by alternative methods. We report extensive comparisons to the common approach of performing enrichment tests between SNP-associated genes and pathway genes. Such comparisons include a new objective method of evaluating pathway association methods, based on the idea that SNP sets for the same disease should yield similar pathways while those for different diseases should yield distinct pathways. Finally, we apply VarSAn to a recently identified set of de novo mutations in a cohort of patients of Hypoplastic Left Heart Syndrome (HLHS) and report on several significant pathways associated with these variants.

## MATERIALS AND METHODS

### Random Walk with Restarts on a heterogeneous network

Random walk with restart (RWR) is a network analysis algorithm that quantifies the proximity of a node to a set of nodes (“query set”) in a graph. We used the RWR implementation in the DRaWR (27) tool as the core of VarSAn. Details of this implementation, especially how it handles a heterogeneous network, are provided in (27). Briefly, given a heterogeneous network with weighted edges, firstly, edge weights are rescaled so that the sum of weights of all edges of the same type is the same constant, regardless of the edge type. (If the network edges are unweighted, all edge weights are assumed to be 1.) Next, transition probabilities are assigned as follows: for any node u, every edge adjacent to it is assigned a transition probability out of u, proportionate to the edge weight so that the sum of transition probabilities out of u equals 1. This step results in a transition probability matrix ***A***, with Aij being the probability that the random walk, if currently at node i, will transition to node j in the next step. (The rescaling step mentioned above ensures that an edge of a more frequent type is less likely to be followed in a transition, all else being equal.) The RWR is a stochastic process where the probability distribution of the random walk’s location at step t, given by ***v**^t^*, evolves as follows:

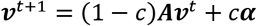

where ‘c’ is the restart probability and ***α*** is the probability distribution over nodes immediately after a restart. We set ***α*** so as to assign zero probability to all nodes outside of the query set and nodes in the query set are assigned equal probabilities summing to 1. Note that the implementation also allows nodes in the query set to have pre-determined node weights, in which case the restart probability at each such node is proportional to its weight. VarSAn uses the node weight feature in the SNP-weight mode. Note that in setting up the heterogeneous network, VarSAn limits the SNP nodes to only include the SNPs in the query set.

The DRaWR program (27) makes a distinction between “gene nodes” and “property nodes”. DRaWR reports the equilibrium probability scores of “property nodes” based on the query nodes selected from “gene nodes”. In using this program in VarSAn we set SNPs and genes as “gene nodes” and pathways as “property nodes”, so that pathways are ranked based on their connectivity to SNP nodes in the query set. SNP-gene edges were added as directed edges with weights of 1 or the appropriate value inferred in the pan-tissue mode.

### Heterogeneous network in VarSAn

Nodes: The network includes SNP nodes representing the query set, gene nodes representing all genes from GTEX, HumanNet and pathway dataset (REACTOME or KEGG), and pathway nodes representing a selection of 321 KEGG (17) pathways or 476 REACTOME (28) pathways. REACTOME pathways were downloaded from https://reactome.org/ and filtered to remove redundancies and small (< 30 gene) pathways (see **Supplementary Note S1, Table S4**). KEGG pathways were obtained from ConsensusPathDB (29).

Coding SNP-gene edges: The Polymorphism Phenotyping v2 (PolyPhen-2) software (26) predicts the functional effect of human non-synonymous SNPs. A (node for a) SNP located within a gene’s coding sequence is connected to (the node for) that gene if it is predicted by PolyPhen-2 to be “probably damaging” or “possibly damaging”. The user may choose to limit connections to “probably damaging” SNPs only.

Non-coding SNP-gene edges: SNP-gene connections for non-coding SNPs (ncSNPs) are based on eQTLs from GTEx analysis v8 (30,31) and/or genomic proximity (distance from gene start site below a user-specified threshold, set to 10 Kbp by default). If the user chooses to define these based on eQTLs, they may further specify the most relevant of 49 tissues for which GTEx eQTLs are available. (The number of SNP-gene associations for each tissue range from 1.1 million to 37 million.) Alternatively, the user may choose to base ncSNP-gene edges on the union of eQTLs across all tissues. (This is the “pan-tissue” option for setting ncSNP-gene connections.) In this case, a SNP-gene edge is weighted by an automatically calculated “relevance score” of the tissue for which the eQTL relationship of the SNP-gene pair has been recorded. The tissue relevance score is calculated as −log(p), where p is the p-value of the hypergeometric test between the query set SNPs and GTEx eQTLs for that tissue. In cases where a SNP-gene pair is an eQTL relationship in multiple tissues, the sum of the appropriate tissue relevance scores is used as the edge weight.

SNP-weight scheme: In this setting, every ncSNP connected to the same gene is first assigned a weight of 1/N, where N is the number of such ncSNPs. Next, the total weight of each ncSNP is set to the sum of weights it is assigned based on all the genes it is connected to (**Supplementary Figure S1**). Every coding SNP is given a weight as 1 in this scheme. This optional scheme allows the user to down-weight the impact of ncSNPs that are in linkage disequilibrium (LD) with each other, as is likely to be the case when multiple ncSNPs connect to the same gene based on eQTL and/or proximity. However, this is a simplistic strategy to deal with the LD issue, motivated by ease-of-use considerations.

Gene-gene and gene-pathway connections: Gene-gene relationships from HumanNet PPI network (32) may be optionally included as edges in the network. These edges were downloaded from (33). Genes belong to a pathway are represented as gene-pathway edges.

### Empirical p-value calculation

VarSAn randomly samples query sets using stratified sampling based on the numbers of coding and non-coding variants in the real query set. Each random query set has the same number of SNPs and the same proportion of coding and non-coding SNPs as the real query set. The empirical p-value of pathway i is calculated as 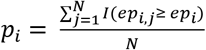, where *ep_i_* is the equilibrium probability of pathway i using the real query set, *eq_i,j_* is the equilibrium probability of pathway i using jth random query set, *I*(*x*) = 1 if *x* is true and = 0 otherwise, N is the number of random query sets (N=100 by default). VarSAn reports all pathway nodes ranked first by empirical p-values and then by equilibrium probabilities.

### Query Sets

Cancer/disease GWAS SNPs: Disease-associated GWAS SNPs were downloaded from GWAS Catalog (34). Breast cancer GWAS SNPs from Michailidou et. al. (35) and prostate cancer GWAS SNPs GWAS Catalog (EFO_0001663) were used. The 12 diseases that have the most associated GWAS SNPs are used in the consistency and discrimination evaluation (see **Supplementary Table S2**).

Cancer somatic mutations: Somatic mutations of breast cancer and prostate cancer were obtained from COSMIC (36). Coding mutations from COSMIC mutation data (genome screens) and non-coding mutations from COSMIC non-coding variants were used.

Drug response GWAS: Drug response GWAS p-values were calculated for each of 24 different drugs/treatments using 284 lymphoblastoid cell lines (37), using the procedure described in (38).

## RESULTS

### VarSAn: A network-guided tool for associating a variant set with pathways

VarSAn is a variant set analysis tool that employs a random walk with restart (RWR) algorithm on a heterogeneous network to rank pathways by their relevance to a given set of SNPs. (We use SNPs and variants interchangeably, and somewhat loosely, to refer to single nucleotide positions that are of interest to the researcher.) The network analyzed by VarSAn must include nodes for SNPs, genes and pathways, as well as edges connecting SNPs to genes and those connecting genes to pathways (**Figure 1**). It may optionally include additional types of nodes and edges. Relevance of a pathway to the given set of SNPs is quantified by the proximity of the pathway node to nodes representing the SNP set, as per the RWR algorithm.

**Figure 1.**
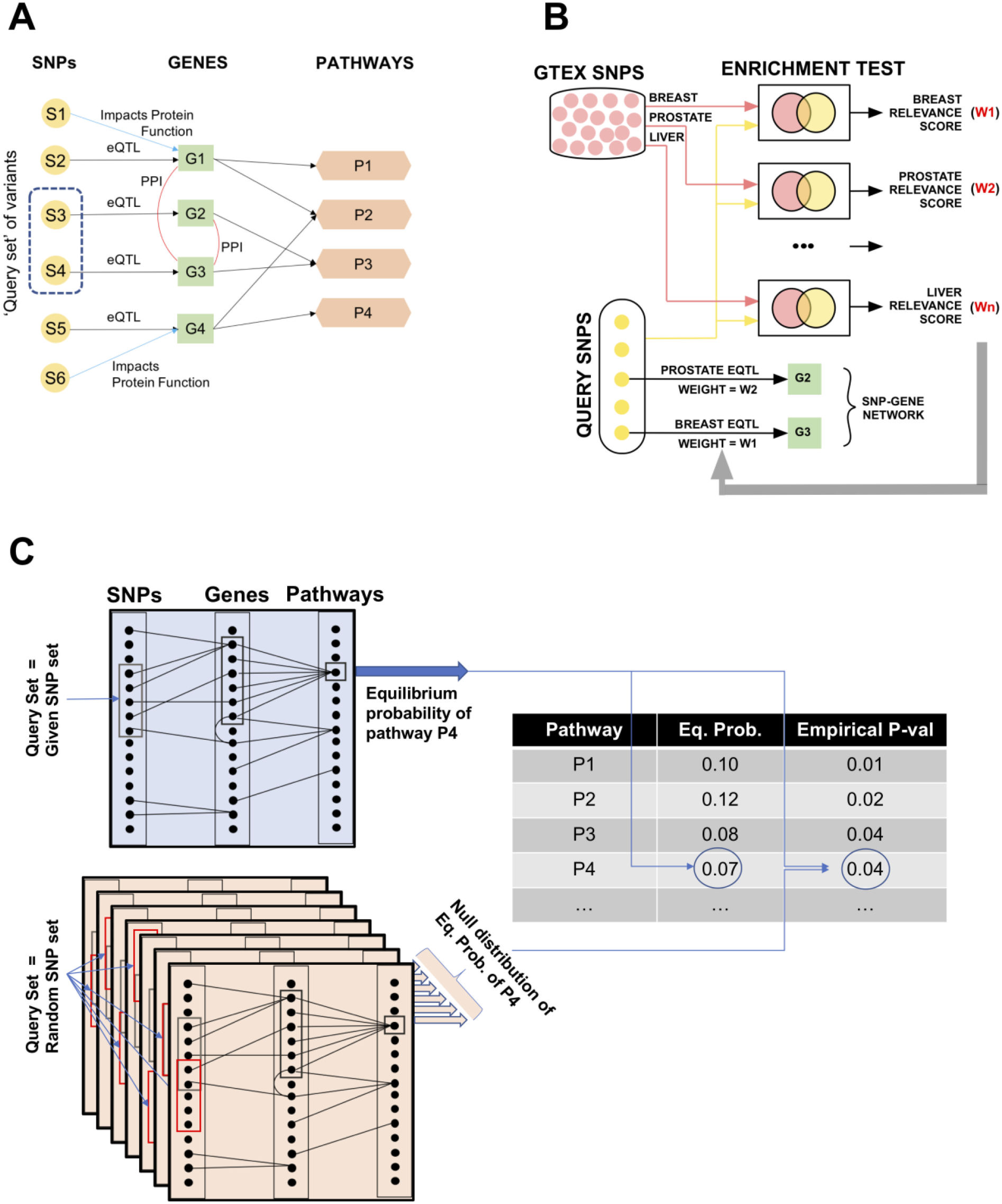
Workflow of variant set characterization by VarSAn. A network is defined with SNP nodes, gene nodes and pathway nodes. SNP and gene nodes are connected by edges representing biological evidence of their relationship, e.g., the SNP is predicted to impact the gene’s encoded protein or the gene’s expression (eQTL). Gene nodes are connected to pathway nodes to indicate genes that belong to a pathway. Gene-gene edges are optional and may represent various types of relationships, e.g., protein-protein interactions. A subset of SNPs, to be characterized, is designated as the “query set” and a Random Walk with Restarts (RWR) algorithm is used, with this query set as its “restart set”, to identify relevant pathways. (B) Empirical p-values reflect the relevance of pathways to the query set of SNPs. Each pathway is assigned an “equilibrium probability” by the RWR algorithm, run with the query set as the restart set. The process is repeated many times, using different random sets of SNPs as the query set, and the equilibrium probability of a pathway from the original RWR is compared to corresponding values with the random query sets to assign an empirical p-value to the pathway. (C) Assignment of SNP-gene edges in the network in pan-tissue eQTL mode. Each SNP-gene edge represents an eQTL in any tissue in the GTEx compendium. The edge is weighted by the “relevance score” (W1, W2, etc.) of the tissue which is calculated as −log(p), where p is the p-value of the hypergeometric test between the query set SNPs and GTEx eQTLs for that tissue. If a SNP-gene relationship is supported by eQTLs detected in multiple tissues, the edge weight is the sum of the relevance scores of those tissues.

The network can be easily customized by the user if so desired (details below), and we present here the general guidelines and our specific decisions in setting up the network for this work. The complete set of SNP nodes in the network (the “SNP universe”) represents a collection of SNPs that includes the user-specified SNP set (the “query set”) as well as other SNPs that are used as a statistical “background” set for contrasting with the query set. Gene nodes represent genes that are related to SNPs in the SNP universe in some pre-determined manner, with those pairwise SNP-gene relationships being represented as edges between respective gene and SNP nodes. Each pathway node represents a biological pathway and is connected to nodes representing genes that belong to the pathway. More generally, pathway nodes may represent any systems-level property or annotation, such as a Gene Ontology (GO) term, shared by a set of genes. Optionally, there may be gene-gene edges representing known relationships, e.g., genetic interactions, among genes. The network allows for multiple types of edges connecting the same two node types, e.g., there may be two types of SNP-gene edges, one representing cases of a SNP predicted to impact a gene’s expression and another representing predicted SNP impact on the activity of the protein encoded by the gene.

Here, we defined one type of SNP-gene edges based on Polyphen2 (26) predictions of coding SNPs impacting protein activity (see Methods). Another class of SNP-gene edges was defined for non-coding SNPs, where we used either eQTL studies to relate SNPs to genes (see Methods) or simply the location of the SNP relative to a gene. We defined gene-gene edges to represent protein-protein interactions from the HumanNet (32) database. Pathway nodes and gene-pathway edges were defined to reflect gene memberships in selected pathways in pathway databases, such as ReactomeDB (28) and KEGG (17) (see Methods).

Once the network and the query set of SNPs has been specified, VarSAn analyzes the dynamics of a random walk with restarts (RWR) whose “restart set” is the query set. Briefly, this simulates the steps of a random “walker” that hops from one node to another over time, at each time step following one of the edges adjacent to the node it currently occupies, with the probability of following an edge being determined by the number and weights of available edges (see Methods). This probability is identical among all edges of the same type, but differs between edges of different types, with rarer edges receiving high relative probabilities to accord them greater weight. In addition to the repeated iterations of such probabilistic hops from node to node, the random walker occasionally (and also stochastically) decides to “restart”, whereby it hops directly to one of the nodes in the restart set regardless of whether such a node is directly connected to the current node. Such a random walk with restarts is well-studied mathematically (39); the “equilibrium probability” of any node reflects the probability of finding the random walker at that node at an arbitrary step, in the long term. Furthermore, due to the occasional but repeated restart decisions by the walker, nodes in, adjacent to or at a few hops from the restart set tend to have higher equilibrium probability, which therefore serves as an objective measure of the network-based proximity of the node to the restart set. VarSAn therefore uses the equilibrium probability of a pathway node as a relative measure of that pathway’s relevance to the query set of SNPs.

#### Accounting for large pathways

The above approach for scoring network nodes tends to be biased toward high-degree nodes in the network. In our case, nodes representing large pathways (with many gene-pathway edges leading into the pathway node) tend to have higher equilibrium probabilities, since there are relatively many ways for the walker to reach such nodes. To address this high-degree node bias, VarSAn calculates “empirical p-values” for every node’s equilibrium probability: it notes the equilibrium probability *p_v_* of a pathway node *v*, then repeats the RWR calculations *N* (say, 100) times, each time using a randomly selected set of nodes as the query set (having the same size as the original query set), counts the number of times (say *n)* out of these *N* repeats that the equilibrium probability of node *v* is greater than *p_v_*, and defines the fraction *n/N* as the “empirical p-value” of node *v* (**Figure 1B**). Note that a node that was assigned a high equilibrium probability mainly due to its large degree (i.e., a large pathway) will tend to have high equilibrium probabilities in the repeats with random query sets as well, for the same reason, and thus have a high (poor) empirical p-value. The empirical p-value tests the significance of a (large) equilibrium probability assigned to a pathway under the null hypothesis that connectivity from the query SNP set to the pathway is no different from connectivity of a random SNP set to this pathway. VarSAn thus ranks pathways first by their empirical p-values, which represent their connectivity to the query set, and then (to break ties) by their equilibrium probabilities, which are determined by both the connectivity and the network topology.

#### Handling multiple SNPs related to the same gene

RWR-based scoring of pathways treats each SNP in the query set as equally important, since the random walker “restarts” by hopping back to one of those SNPs chosen uniformly at random. If multiple SNPs in the query set are related to the same gene, then that gene will have greater influence on pathway scoring compared to a gene that is related to a single SNP in the query set. This may or may not be desirable to the user. For instance, consider a case where a GWAS-based query set includes multiple non-coding SNPs that are in linkage disequilibrium (LD) with each other and are located in the regulatory region of the same gene. In this case, the RWR approach assigns relatively high weight to the gene, and thus prefers pathways that include the gene. However, such proportional weighting may be undesirable, since the multiple SNPs in LD do not provide independent evidence for the significance of that gene (and its related pathways) to the disease represented by the query set. On the other hand, proportional weighting may be deemed appropriate by the user, if for instance the query set was derived by means other than association tests. and multiple SNPs near the same gene do in fact present greater evidence for that gene’s relevance. The VarSAn implementation offers the user two simple modes for handling of multiple non-coding SNPs related to the same gene. The first mode (called “SNP-weight”, see **Supplementary Figure S1**) is to assign each of *N* non-coding SNPs connected to the same gene a weight of 1/*N*; the RWR algorithm uses these weights during the restart step, hopping back to a SNP node with probability proportional to its weight. (Note that a SNP node may be connected to multiple genes, in such cases its weight is the sum of its assigned weights due to its connection to each gene.) In the other mode (called “SNP-uniform”), all SNPs have equal weight. Note that this dichotomous handling of SNPs connected to the same gene applies only to non-coding SNPs (see Discussion).

### Case studies with VarSAn: pathways related to breast cancer and prostate cancer

As a first assessment of its functionality, we used VarSAn to identify pathways related to breast cancer GWAS SNPs, with the hope of cross-checking the top reported pathways against prior knowledge about this disease. The SNP set, obtained from GWAS Catalog (35), includes 137 non-coding SNPs and 9 coding SNPs. VarSAn was run with a network defined as above, with SNP-gene edges based on eQTLs (located within 1 mbp of gene) for breast tissue from the GTEx project (30,31) (for non-coding SNPs) or PolyPhen-2 predictions of “probably damaging” and “possibly damaging” SNPs (for coding SNPs). **Table 1A** lists the 15 top-ranked pathways (among 476 pathways in the network) from this analysis; for 12 of these pathways, the table cites a prior publication supporting the pathway’s relevance to breast cancer (also see **Supplementary Table S1**). For example, the pathway “PKMTs_methylate_histone_lysines” is ranked second: lysine methylation and demethylation are catalyzed by protein lysine methyltransferases (PKMTs), a process shown to be important in cancers, including breast cancer (40). The third ranked pathway – “Hedgehog_on_state” – points to the involvement of Hedgehog signaling, which is known to mediate breast cancer invasiveness (41). Three of the top pathways identified (**Table 1A**) are p53-related; the tumor suppressor p53 is found to be altered in 20%-40% of breast carcinomas cases (42), and p53-related pathways are among the most intensely studied in this disease context (43). The pathway “Stabilizaton of p53” was among the highly ranked pathways (ranked 7, **Table 1A**), and provides a useful glimpse into the inner workings of VarSAn. This pathway contains 73 genes, yet only one of those genes is related directly to the query set of SNPs (GWAS SNP chr22_28725099 is an eQTL of the checkpoint kinase gene CHEK2), which would normally be insufficient evidence of the relevance of this pathway. However, as **Figure 2A** illustrates, VarSAn not only uses the corresponding “two-hop path” (SNP to gene to pathway), it also finds 27 “three-hop paths” (SNP to gene to interacting gene to pathway) and is thus able to relate as many as 26 of the 146 SNPs in the query set to the “Stabilization of p53” pathway, thereby ranking the latter as highly relevant to the query set.

**Table 1.**
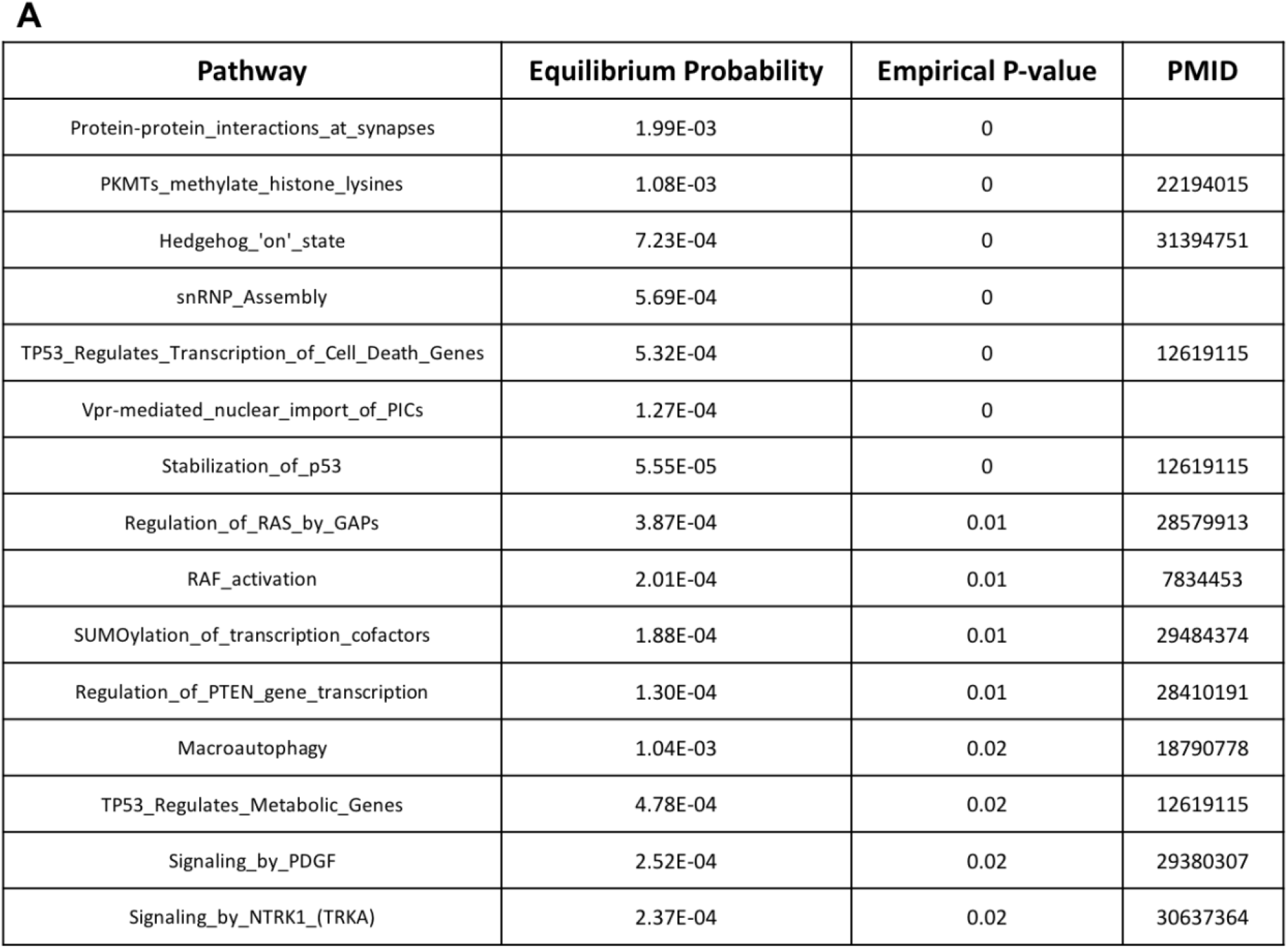
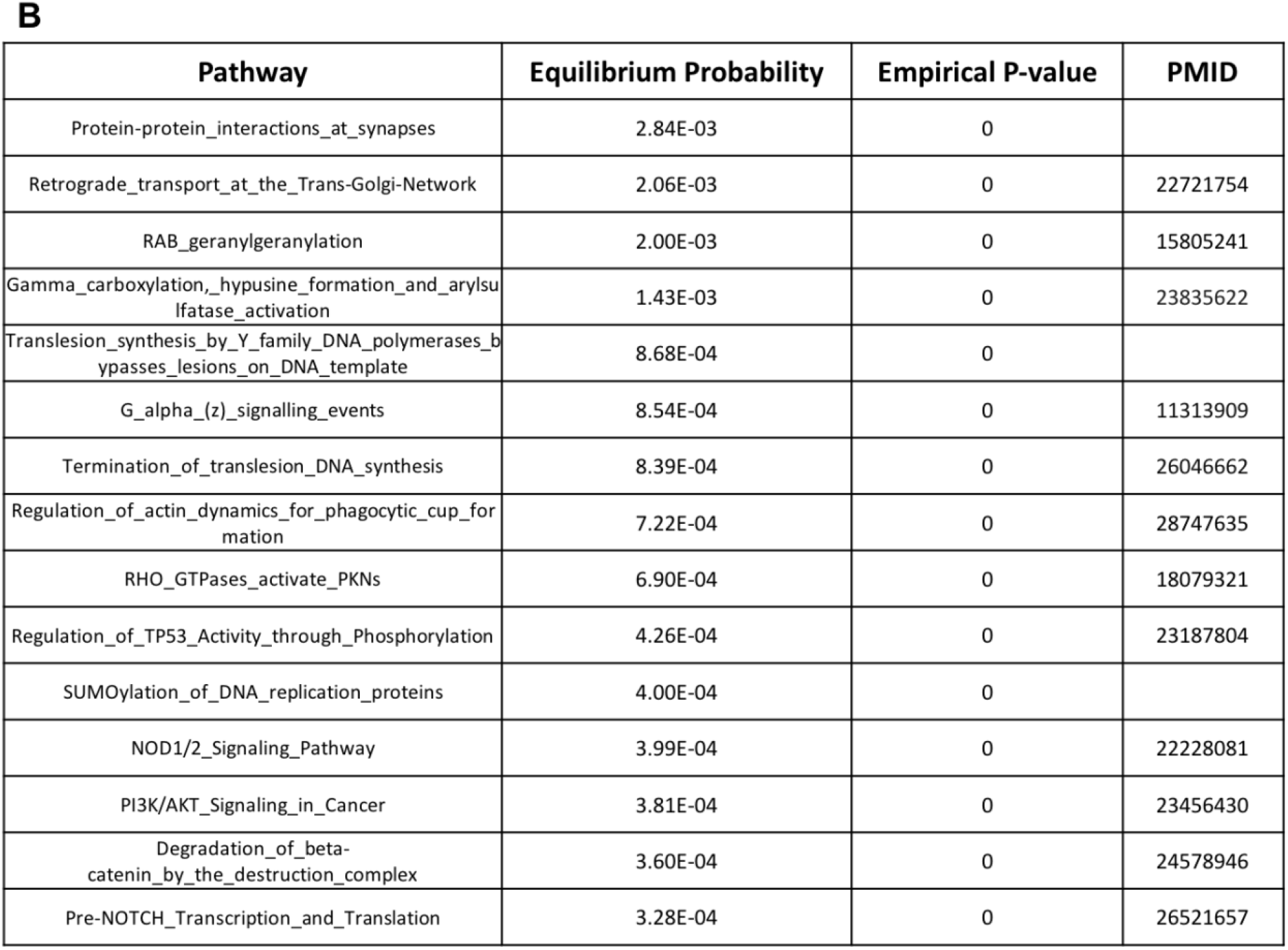
Top 15 pathways reported by VarSAn for a query set of GWAS SNPs for (A) breast cancer and (B) prostate cancer, along with citations to literature evidence supporting their relevance to the respective disease. Pathways are ranked first by empirical p-value and then by equilibrium probability.

**Figure 2.**
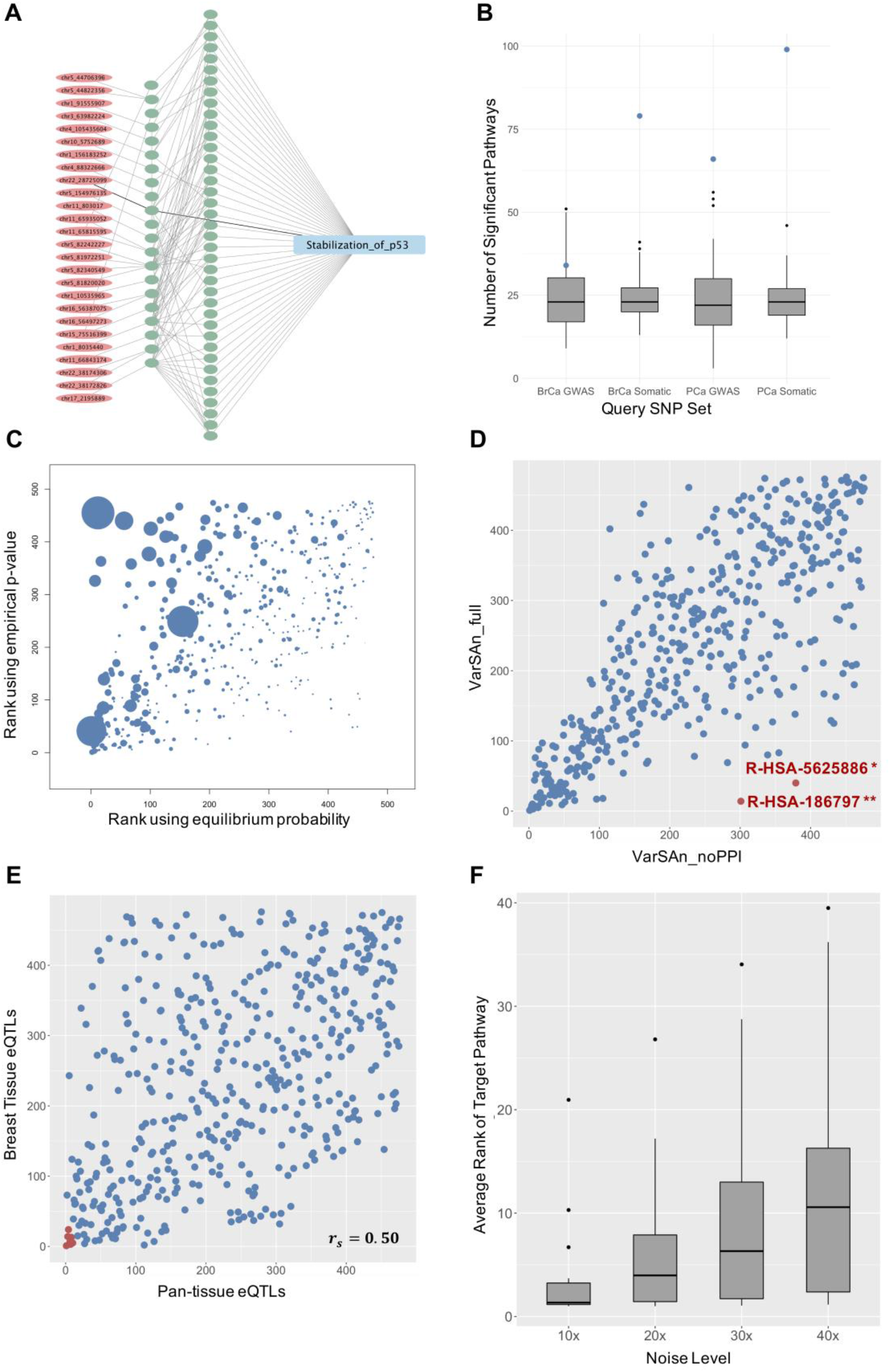
A) SNP (red) and gene (green) nodes connected to the pathway node “Stablization_of_p53”, directly (SNP – gene – pathway) or indirectly (SNP – gene – gene – pathway) are shown. The left layer of gene nodes are genes directly connected to SNP nodes. The SNP set, obtained from a breast cancer GWAS study, has only one SNP that is associated with a member gene of the pathway (darker edge), but many indirect connections establish the relevance of this pathway to the SNP set. B) Numbers of significant pathways (at empirical p-value <= 0.05) reported by VarSAn for each of four disease-related SNP sets (blue dots). Each number is contrasted with a distribution (box and whiskers plots) of corresponding numbers for 100 random query sets of similar size and composition as the disease-related SNP set. C) Pathway ranks reported by VarSAn for the BrCa GWAS SNP set, when using equilibrium probability (x-axis) or empirical p-value (y-axis) as pathway score. Size of each circle represents the number of genes in the pathway. D) Effect of using gene-gene edges. Pathways ranked highly (top 50) by one method and ranked at 250 or worse by the other method are marked in red. * Reactome pathway R-HSA-5625886: Activated PKN1 stimulates transcription of AR regulated genes KLK2 and KLK3; ** Reactome pathway R-HSA-186797: Signaling by PDGF E) Scatter plot of pathway rank using VarSAn on the BrCa GWAS SNP set, with SNP-gene edges based on eQTLs from the relevant tissue (y-axis) versus those based on eQTLs from all tissues (pan-tissue approach, x-axis). Seven of the top 10 pathways reported by the pan-tissue approach are in the top 50 reported with the breast tissue eQTLs being used for SNP-gene edges (marked in red) F) Rank of target pathways for 24 drugs with 10x, 20x, 30x and 40x noisy query sets. Each box represents average rank of target pathways across the 20 independent tests at corresponding noise level.

We performed a similar subjective assessment of VarSAn on a set of 57 GWAS SNPs (2 coding, 55 non-coding) for prostate cancer (35), cross-checking the top reported pathways against the literature (**Table 1B**). The pathway “RAB_geranylgeranylation” was ranked 2 among 476 pathways. Many RAB genes have been found up- or down-regulated in prostate cancer. For example, increased RAB25 (44,45) levels have been observed in prostate cancer and Rab3B (46) has been noted to promoter cancer cell survival. As another example, the “NOD1/2 Signaling Pathway” (rank 12 in **Table 1B**) has been shown to play an important role in prostate cancer progression (47). We also used VarSAn with query sets comprised of somatic mutations found in breast cancer or prostate cancer patients from the Cosmic database (36). While the GWAS-based query sets analyzed above included 50-150 SNPs each, with over 90% being non-coding SNPs, the somatic mutation-based query sets are much larger and are more biased towards coding variants (16,011 coding variants and 3,005 non-coding variants in breast cancer; 11,449 coding and 1,365 non-coding variants in prostate cancer). These two query sets thus exhibited different statistical properties while still representing the same two disease contexts as above. Top pathways reported by VarSAn are shown in **Supplementary Table S1**, and include several pathways known to be associated with the corresponding cancer type.

Results presented above illustrate that VarSAn can identify biologically relevant pathways for SNP sets of diverse sizes and composition (coding versus non-coding). Next, as a purely statistical assessment of the results, we asked how the number of significant pathways discovered in each of the four analyses above compares to random expectation. To this end, we generated a random query set with the same number of SNPs and same proportion of non-coding and coding SNPs as the original query set, and counted the number of significant pathways (with empirical p-value <= 0.05). We repeated this 100 times for each of the four disease-related query sets introduced above. **Figure 2B** shows the number of significant pathways observed in these randomized “control experiments”, with the blue dots marking the number of significant pathways VarSAn reports for the respective cancer-related query sets. Approximately 24 out of 476 pathways are expected to be significant by chance at p <= 0.05. Indeed, most of the random query sets were reported as related to 20-30 significant pathways, small in comparison to corresponding numbers for cancer-related query sets (PCa somatic mutations: 99, BrCa somatic mutations: 79, PCa GWAS: 66, BrCa GWAS: 34).

### Evaluation of methodological choices in VarSAn

We next assessed the value of some of the key methodological choices made in implementing VarSAn. For instance, as noted above, VarSAn not only calculates equilibrium probabilities of pathway nodes (representing network connectivity of each node to the query set), it then computes empirical p-values of those equilibrium probabilities. This is expected to mitigate the well-known tendency of RWR algorithms to assign greater probability to high-degree nodes (large pathways in our case). **Figure 2C** compares the ranks of all pathways related to the BrCa GWAS query set, based on equilibrium probabilities (x axis) and on empirical p-values (y axis). We observe that large pathways tend to be ranked near the top when only equilibrium probabilities are used while they are more evenly distributed when empirical p-values are considered, thus arguing that VarSAn addresses the bias towards large pathways.

A second key feature of VarSAn is the inclusion of gene-gene relationships in the network. (This is an optional feature and in our tests above we used a protein-protein interaction database to define such edges.) This allows indirect SNP-to-pathway connections to contribute to pathways’ equilibrium probabilities, as we saw through an example (“Stabilization of p53” pathway associated with BrCa GWAS set) above. While direct connections capture the extent to which genes related to the given SNP set *belong to* a pathway, indirect connections relax this view to accommodate instances of SNP-related genes being “partners” of pathway genes; however, such indirect connections contribute less to VarSAn’s calculations compared to direct connections (see Methods). **Figure 2D** compares the ranks of pathways from VarSAn analysis of the BrCa GWAS query set, with and without gene-gene edges in the network. (We will refer to these two settings of VarSAn as VarSAn_full and VarSAn_noPPI.) There is a strong agreement between the two rankings, but there are also conspicuous examples of pathways highly ranked by VarSAn_full and poorly ranked by VarSAn_noPPI. For instance, the pathways “Signaling by PDGF” (54 genes, rank 14) and “Activated PKN1 stimulates transcription of Androgen Receptor regulated genes KLK2 and KLK3” (36 genes, rank 40) were ranked at 250 or worse by VarSAn_noPPI. We have already noted the literature-based relevance of the PDGF signaling pathway (**Table 1**). The role of Androgen Receptor-controlled expression of genes klk2 and klk3, which encode human kallikrein 2 (hK2) and prostate specific antigen (PSA) respectively, is well-documented for prostate cancer, where these genes are used as biomarkers. Recent studies have yielded promising new insights into their diagnostic/therapeutic potential for BrCa (48), and the broader role of Androgen Receptor in BrCa is a topic of active research today (49). Thus, in light of literature evidence, these two pathways revealed by VarSAn as relevant to breast cancer, but only when using gene-gene interactions, illustrate how allowing for indirect connections between SNPs and pathways can provide novel biological insights. (We also noted in **Figure 2D** that there are no pathways with a similar disparity of rankings in the opposite direction, i.e., a pathway highly ranked, say <= 50, by VarSAn_noPPI that is poorly ranked, say >= 250, by VarSAn_full.)

We next assessed the impact of methodological choices made in defining SNP-gene edges in the network. It was noted above that a non-coding SNP is deemed related to a gene if it is an eQTL of and located in cis-regulatory region (1 Mbp up- and down-stream of the transcription start site) of the gene. In our analyses above, for each cancer (breast or prostate) we relied on eQTLs detected by the GTEx project (30,31) for the corresponding tissue-of-origin in defining SNP-gene edges. On the other hand, if the user-specified SNP set does not have an obvious correspondence to a tissue for which a comprehensive collection of eQTLs exists, VarSAn adopts the following strategy (**Figure 1C**: first, it considers GTEx eQTLs for every tissue separately; second it assigns a weight to each tissue that reflects how enriched the query set of SNPs is for eQTLs from that tissue, thereby scoring the relevance of each tissue to the query set; and finally, it creates a SNP-gene edge in the network if the SNP is an GTEx eQTL of the gene in any tissue, but weights the edge based on the tissue-relevance computed in the previous step (see Methods). Thus, if the query set comprises breast cancer-related SNPs, it is likely that breast tissue will be scored highly, and SNP-gene edges based on breast tissue eQTLs will receive high weights. The ability to relate SNPs to genes automatically by utilizing a pan-tissue compendium of eQTLs is an important feature of VarSAn that enables its broader applicability. Here, we sought to determine how the results of adopting this pragmatic strategy differ from the ideal scenario where eQTL data from the appropriate tissue exist. For this, we compared pathway rankings for the BrCa GWAS query set, obtained as above using breast tissue eQTLs, with rankings produced by using the less specific, pan-tissue approach to SNP-gene relationships. As shown in **Figure 2E**, the two rankings are highly correlated (Spearman’s rho 0.50, p-value 1.45E-31). For example, of the top 10 pathways reported by the pan-tissue approach, seven are in the top 50 reported with the breast tissue eQTLs being used for SNP-gene edges. A high correlation was observed also when repeating the above assessment using the PCa GWAS query set (**Supplementary Figure S2**). These results suggest that the strategy of utilizing pan-tissue information on SNP-gene relationships in creating the VarSAn network yields pathway rankings that are reasonable approximations to those based on a more query-specific tissue. Finally, we compared the default strategy (VarSAn_full) to a baseline where SNP-gene edges were based solely on location (10 kbp up- and down-stream of the gene), and noted that the pathway rankings are very different between the two strategies (**Supplementary Figure S3**).

### Robustness of VarSAn to irrelevant SNPs in query set

In our experiences with the VarSAn tool, we noticed that the top pathway associations were often based on (SNP – gene – pathway) connections involving only a small fraction of the SNPs (variants) being analyzed. This is a desirable property: the variant set is likely to include only a relatively small number of variants easily linked to a particular pathway, and the variant set characterization method should be able to recover that pathway despite the presence of a large number of “irrelevant” variants in the query set. To test this aspect of the VarSAn tool, we set up a controlled analysis scenario where we started with a small set of SNPs with a strong connection to a specific pathway *P* (this ensured that VarSAn ranks the pathway *P* at or near the top), and progressively introduced many randomly chosen variants to the query set, examining how the rank of the pathway *P* changes in these more “noisy” conditions.

For this analysis, we utilized genotype, gene expression and drug response data on a panel of ~300 lymphoblastoid cell lines (LCLs) from a prior study (38). This data set includes measurements of cytotoxicity (*EC*_50_ of dosage-response curve) of each of of 24 different treatments, mostly cancer drugs, on every LCL. We used genotype data from the LCL panel to define GWAS SNPs for each treatment (using cytotoxic response as phenotype); while there were few or no SNPs that met standard GWAS thresholds (after multiple hypothesis correction), we used the 100 most significant p-values to define the GWAS SNP set for that treatment. For each drug, we first used VarSAn to associate the 100 most significant SNPs with REACTOME pathways, and selected the top-ranked pathway as the “target” pathway. We then use VarSAn to identify the top 5 SNPs with strongest connections to the target pathway (for each drug). The resulting set of 5 SNPs was considered the “clean query set”, and its association with the target pathway was considered as “true” (by design). We then added 50, 100, 150, or 200 randomly selected SNPs to the query set to construct “noisy query sets” with noise levels of 10x, 20x, 30x and 40x respectively. Twenty noisy query sets were generated independently for each noise level. We noted down the rank of the target pathway among all 476 REACTOME pathways, upon applying VarSAn to the noisy query sets, and noted the average rank across the 20 independent tests at each noise level (**Supplementary Table S3**). We repeated this entire process for each of the 24 drugs. The results, shown in **Figure 2F**, indicate how VarSAn retains or loses its sensitivity to the true association with noise level. We noted that on average (across all drugs), the true pathway continues to be reported in the top 5 even at the 20x noise level (20 times as many “irrelevant” SNPs as the clean query set), and in the top 10 at the 30x noise level. At the 40x noise level the true pathway is no longer in the top 10, on average. However, we also noted that the above behavior varies substantially from one drug to another, implying that the strength of connection between the clean query set and the target pathway can vary substantially, and in some cases the association remains discoverable (e.g., average rank of 5 or better) even at the highest noise level applied. Overall, this exercise suggest that VarSAn can effectively recover the pathway associated with a small set of variants even in the presence of tens of times as many irrelevant variants in the query set.

### Comparison of VarSAn with overlap-based statistical tests

The conventional approach to associating SNP sets with pathways is to find the genes related to the given SNPs and then test for associations between the resulting gene set and each pathway, using, say, the Hypergeometric test. We thus compared the results of VarSAn to those from such an approach, which we refer to as the “HGT” method. In this method, the query SNP set is mapped to a gene set using the same rules as in VarSAn, and the gene set is tested for enrichment in genes of a pathway using the Hypergeometric test. The latter step is repeated for each pathway, and pathways are ranked by p-value. **Figure 3A** compares this ranking with that from VarSAn, for the BrCa GWAS SNP set. Firstly, we note that the results are in broad agreement. For instance, the pathways have a strong tendency to be ranked <= 100 by both methods or >= 100 by both methods. (422 of the 476 pathways are in these two square regions of the plot, compared to 318 by random chance.) Secondly, we focused on pathways with highly disparate rankings between methods, specifically, a rank <= 50 by one method and >= 250 by the other. There are only six such pathways, and five of them are highly ranked by VarSAn (but poorly ranked by HGT). The latter five pathways have literature evidence supporting their role in BrCa (**Table 2A**), suggesting to us that VarSAn’s network-based approach can identify important pathways that are not discovered using the conventional approach. There is a single pathway, Interferon gamma signaling, that was highly ranked by HGT but ranked poorly by VarSAn, reminding us that the two approaches can provide complementary insights.

**Figure 3.**
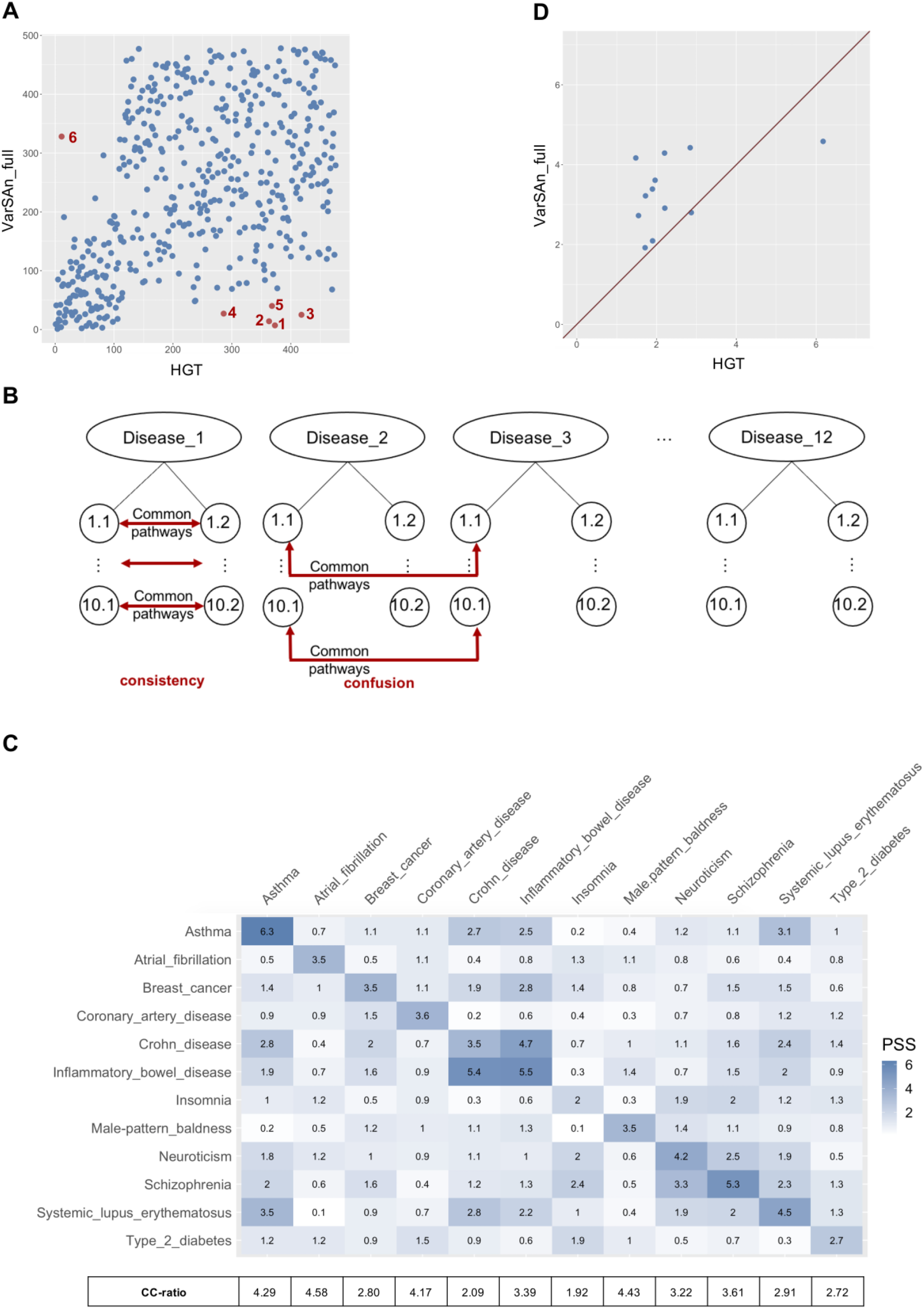
A) Scatter plot of pathway ranks reported by VarSAn and HGT, for the BCa GWAS SNP set. Pathways labeled 1 to 5 are those with significantly better ranking in VarSAn than in HGT, with supporting literature evidence in Table 2A. Pathway labeled 6 is “Interferon gamma signaling”, and was ranked significantly better by HGT. B) Scheme for assessing consistency and confusion scores of a pathway ranking method. SNPs associated with each disease are sampled to create two mutually exclusive subsets of 100 SNPs each; the process is repeated 10 times to create 10 pairs of subsets. Top pathways reported for each of the two subsets in a pair should be similar, and this is captured in the consistency score. Top pathways for SNP sets from different diseases should be distinct, and this is reflected in the confusion score. C) Consistency and confusion scores for all diseases and disease pairs in the evaluation. Diagonal entries represent consistency score of different diseases. Off-diagonal entries represent confusion scores between each pair of diseases. There are two entries for each disease pair, due to an asymmetry in the evaluation procedure (see Methods); these two entries may be considered as two estimates of the confusion score. CC-ratio of each disease is calculated as the ratio of consistency score over the average of all confusion scores where the target disease is a member of the pair. D) Scatter plot of CC-ratios of VarSAn vs HGT.

**Table 2.** A) Five pathways ranked within top 50 by VarSAn but worse than 250 by HGT, for the BCa GWAS SNP set. Citations (Pubmed identifiers) provide literature evidence of their relevance to breast cancer. B) Ranks of a “positive control” pathway for each disease (breast cancer or prostate cancer), obtained from the KEGG database, as per the two methods (VarSAn and HGT). The method was required to rank a large collection of 321 KEGG pathways for relevance to either a GWAS set or a somatic mutations set for the respective disease.

**A**
|  | Pathway | PMID |
| --- | --- | --- |
| 1 | Stabilization_of_p53 | 12619115 |
| 2 | Signaling_by_PDGF | 29380307 |
| 3 | Ubiquitin_Mediated_Degradation_of_Phosphorylated_Cdc25A | 10995786 |
| 4 | PRC2_methylates_histones_and_DNA | 23251464 |
| 5 | Activated_PKN1_stimulates_transcription_of_AR_(androgen_receptor)_regulated_genes_KLK2_and_KLK3 | 10188912 |

**B**
| Query Set | VarSAn | HGT |
| --- | --- | --- |
| Prostate Cancer (somatic) | 40 (0) | 55 (0.91) |
| Prostate Cancer (GWAS) | 32 (0.02) | 47 (0.59) |
| Breast Cancer (somatic) | 56 (0.01) | 246 (0.99) |
| Breast Cancer (GWAS) | 104 (0.32) | 74 (0.97) *** |

The KEGG pathway database (17) includes disease-related pathways including a “breast cancer pathway” and a “prostate cancer pathway”, which offered us another opportunity to compare the two ranking methods (VarSAn and HGT). For this evaluation, we used 321 KEGG pathways to define pathway nodes in the network (evaluations above were based on REACTOME pathways), and used VarSAn and HGT respectively to rank these pathways for relevance to BrCa and PCa GWAS and somatic mutation query sets (**Table 2B**). For three of the four test cases, VarSAn ranked the known disease-related pathway higher (for a query set representing that disease) than did HGT, with the most notable difference being for the BrCa somatic mutation SNP set, where the breast cancer pathway was ranked at 56 by VarSAn and at 246 by HGT. Moreover, in these three cases, the empirical p-value assigned by VarSAn to the known disease-related pathway was <= 0.05, while the Hypergeometric test p-value was >= 0.5. In one of our four tests (BrCa GWAS query set, **Table 2B**), HGT ranked the known pathway higher than VarSAn, although both methods reported insignificant p-values (0.97 and 0.32 respectively), suggesting that they both failed to detect the pathway’s association with this query set.

### A novel evaluation framework based on consistency and discrimination

Our evaluations above relied on the use of external annotations of pathways relevant to the query set being analyzed, introducing an element of subjectivity to them, since “relevance” is subjective. This is a problem faced by any current framework for testing pathway associations. To address this, we devised a fully data-driven scheme for such evaluations, based on the following basic principle: if we have different sets of SNPs associated with different diseases, a good pathway association method should report the same pathways when analyzing random subsets of SNPs for the same disease (“consistency”) and report different pathways when analyzing SNP sets of different diseases.

To implement this scheme, we first obtained GWAS SNP sets for 12 diseases from the GWAS Catalog (34) (**Supplementary Table S2**), ensuring that there were at least 200 GWAS SNPs for each disease. From each such SNP set, we derived 10 pairs of mutually exclusive subsets of size 100 via random sampling (see Methods, **Figure 3B**). Each of the 20 SNP sets thus defined for each of the 12 diseases was then used as a query set for VarSAn to rank pathways. We defined the “pathway similarity score” (“PSS”) between any two SNP sets as the number of pathways that have a top-20 rank of association with both SNP sets. The PSS was computed between each pair of mutually exclusive subsets of SNPs of the same disease, and its average across 10 such pairs was noted as the “consistency score” of rankings for that disease. Similarly, the PSS was calculated for 10 pairs of SNP subsets representing two different diseases (see Methods), and its average across the 10 pairs was noted as the “confusion score” of pathway rankings for that pair of diseases.

**Figure 3C** shows the consistency scores (diagonal values) and confusion scores (off-diagonals) for VarSAn rankings, across all diseases and disease-pairs respectively. (There are two off-diagonal entries for each disease pair, which represent two separate estimates of the confusion score, see Methods.) We noted that the diagonal entry is almost always the largest in its respective row or column, indicating that the top pathways reported for two (non-overlapping) sets of SNPs for the same disease tend to be more in agreement than pathways lists reported for different diseases. For instance, the consistency score for “Asthma” is 6.3, meaning that on average 6.3 pathways are shared among the top 20 pathways reported from two mutually exclusive sets of GWAS SNPs for this disease. In comparison the confusion score for Asthma and any other disease is about 1.5 on average (mean of off-diagonal entries in first row and first column), revealing that the pathway lists reported for Asthma and one of the other 11 diseases tend to have far less commonality. The relatively few high values outside of diagonals in **Figure 3C** point to pairs of diseases with high pathway-level similarity between their SNP sets. For instance, “Crohn disease” and “Inflammatory bowel disease” have confusion scores similar to their respective consistency scores; this is expected since the two diseases (as defined in the GWAS Catalog) are related (50). Similarly, “Neuroticism” and “Schizophrenia” had high mutual confusion scores (within two-fold of their individual consistency scores); indeed, there is some literature evidence suggesting neuroticism as a risk factor for schizophrenia (51). Note that the GWAS SNP sets representing these two phenotypes do not have a significant overlap (739 Schizophrenia GWAS and 642 Neuroticism GWAS share only 7 common SNPs), and the link reported in **Figure 3C** emerges through sharing of pathways as revealed by the SNP sets. An intriguing off-diagonal entry is for the pair Asthma and “Systemic lupus erythematosus” (SLE), where the confusion score is relatively high and within a factor of two of the individual consistency scores. This reflects a greater than usual extent of shared pathways between these diseases, and may be related to literature evidence of their shared Immunoglobin E (IgE)-related pathophysiology (52) and reports of increased risk of Asthma in SLE patients (53).

However, other than the few above-mentioned interesting examples of larger off-diagonal entries, the matrix in **Figure 3C** clearly exhibits greater consistency scores than confusion scores, indicating that GWAS SNP sets of different diseases lead to different pathway lists. This is a desirable feature of the pathway ranking method, as mentioned above. To make this contrast between consistency and confusion scores more explicit, we calculated, for each disease, the ratio of its consistency score (diagonal value) and the average confusion score against every other disease (average of off-diagonal values in the row and column for that disease). This ratio, called the CC ratio, is shown at the bottom of **Figure 3C**, and has a median of ~3.3. The CC ratio offers us an objective way to evaluate a pathway ranking method and to compare methods as well. We thus calculated consistency and confusion scores for the HGT method, in a manner identical to how **Figure 3C** was derived. The result, shown in **Supplementary Figure S4**, again tend to place large values on the diagonal, but now the off-diagonal values are frequently comparable to diagonal values in the same row or column, suggesting a smaller gap between consistency and confusion scores. To investigate this, we calculated CC ratios for the HGT method (shown at the bottom of **Supplementary Figure S4**) and plotted these along with corresponding ratios for VarSAn in **Figure 3D**. We see that VarSAn has a greater CC ratio for 10 of the 12 diseases, strongly suggesting its advantage over the HGT approach for retrieving pathways related to disease SNPs.

### VarSAn identifies pathways associated with Hypoplastic Left Heart Syndrome

Hypoplastic Left Heart Syndrome (HLHS) is a congenital heart defect where the left side of the heart is critically underdeveloped. It is a rare, heritable disease (54) and despite promising recent findings (55–58) its genetic basis is poorly understood (59) and mortality remains substantial (60). Here, we analyzed sequencing data from 24 trios (HLHS-affected proband and unaffected parents) (61), identified 654 *de novo* single-nucleotide variants from the cohort (**Supplementary Table S5**), and used VarSAn to characterize systems-level processes and pathways implicated by the variant set. Variant-gene edges in the network were determined based on proximity alone (variant located within 10kb up- or downstream of gene) since eQTL information from an appropriate tissue is not available. Gene-gene edges were based on protein-protein interactions from HumanNet database, and the pathway nodes represented REACTOME pathways, as in analyses above.

Twenty eight of the 476 pathways were determined to have empirical p-value < 0.05 (**Supplementary Table S1**), and of these at least ten pathways (**Table 3**) have been implicated in HLHS or heart development in the literature. For instance, the “VEGFA-VEGFR2 Pathway” (empirical p-value 0.02, rank 15) has been studied in the context of HLHS and congenital heart disease (62) and failure to upregulate VEGF noted (63). This association was discovered based on 10 de novo variants that are located near genes of the pathway (**Supplementary Figure S5**) as well as 235 variants near genes that are not in the pathway, but interact with genes of the pathway (**Supplementary Figure S6**). Another interesting association identified was that with the mTOR signaling pathway, which is known to play a role in embryonic cardiovascular development (64); disruptions to this pathway are believed to cause ventricular wall thinning and cardiomyocyte apoptosis (65). These and other examples of pathways associated by VarSAn to HLHS variants provide avenues for future research into the systems biology of this disease.

**Table 3.**
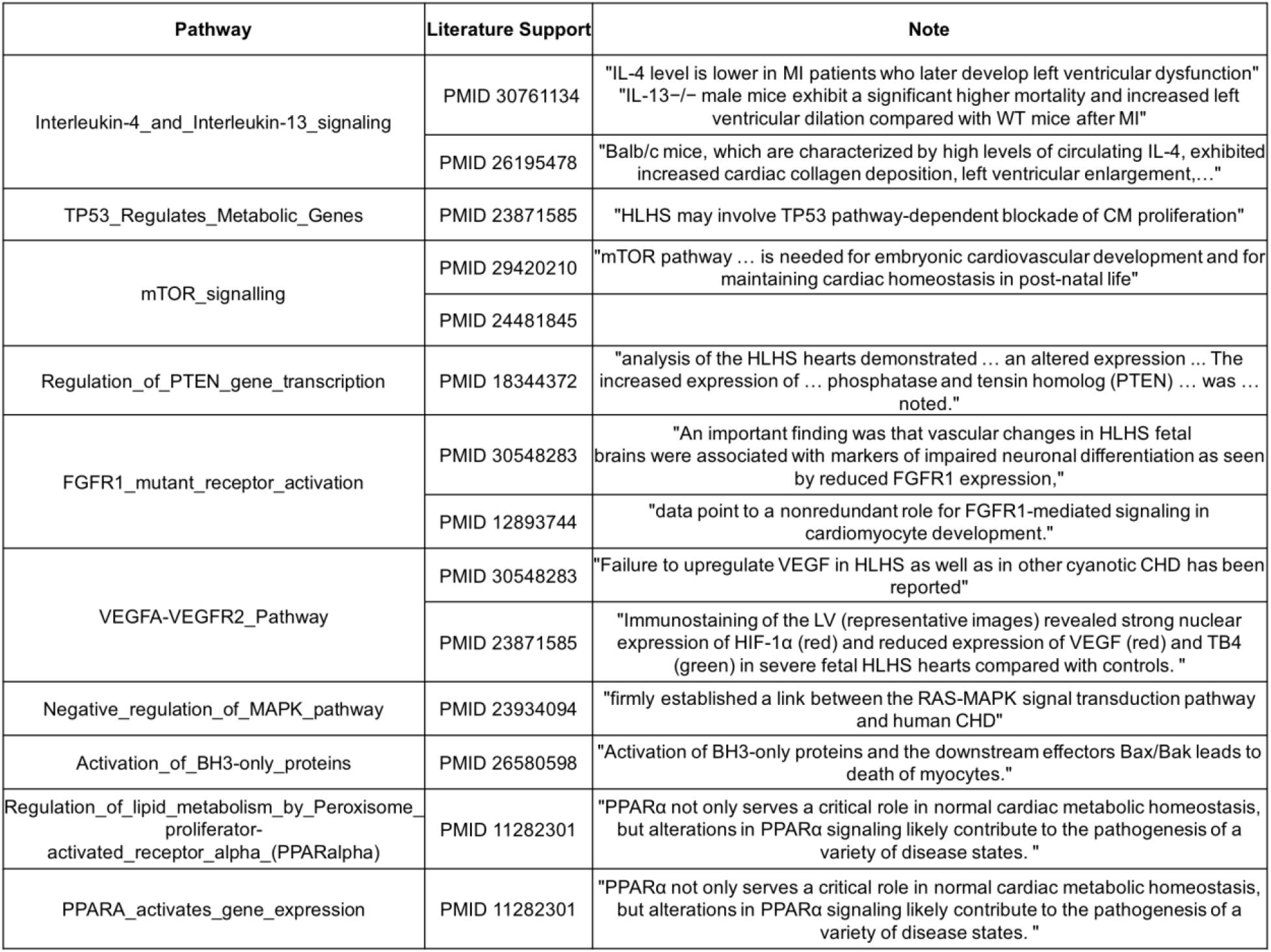
Top-ranked pathways reported by VarSAn for query set of HLHS de novo mutations. Notes are excerpts from cited publications.

## DISCUSSION

Our goal in this work was to develop a tool for characterization of pathways relevant to a given set of SNPs. This is conceptually similar to the task of characterizing pathways associated with a gene set, a well research problem for which several solutions have been proposed (66). One of the more popular approaches for gene set characterization is based on over-representation analysis, exemplified by the Hypergeometric test (HGT) of overlap between a given gene set and pathway genes. The HGT approach has a natural extension to SNP set characterization, whereby the SNP set is mapped to a gene set, which is then tested for pathway association. We tested this approach as a point of comparison to VarSAn, and found objective and subjective evidence that the latter is able to identify biologically interesting pathway associations not revealed by HGT. It is worth considering some of the theoretical differences between the two approaches. Firstly, in the HGT approach, a SNP either “relates to” a pathway or it does not, while VarSAn implements a quantitative notion of SNP relationship to a pathway. In particular, a SNP may be connected to a pathway directly (path via one gene) or indirectly (path via two or more genes), and these two types of connection have different weights. Our case studies illustrated (**Figure 2A**) how indirect connections may lead to meaningful pathway discoveries. Even direct connections (SNP – gene – pathway) may not always have the same weight. For example, a SNP connected to a gene that belongs to exactly one pathway has a stronger pathway connection than a SNP connected to a gene that belongs to several pathways. As another example, a SNP connected to two genes in the same pathway provides quantitatively different information about pathway relevance than if it was connected to only one of those genes. A second difference between VarSAn and HGT is that the former is inherently a pathway ranking scheme where pathways are scored (for relevance) relative to each other, while the latter evaluates each pathway’s relevance by an absolute statistical criterian (significance of overlap). This gives HGT the advantage that the redundancies in the pathway compendium do not affect scoring of individual pathways, although such redundancies naturally translate to redundancies in reported associations and have been noted as a practical concern (67).

It is worth noting that there are alternative methods, e.g., based on the statistics underlying “gene set enrichment statistic” (GSEA), to identify pathways related to GWAS SNPs (15). While GSEA-based approaches have certain advantages compared to a HGT-based approach, their use for GWAS pathway analysis relies on first ranking SNPs based on GWAS p-values, while our focus in this work was on the case where the user seeks to analyze a set of SNPs, without further information on the relative ranking of those SNPs. Such a scenario may arise, for example, from family-based studies of a disease (68). At the same time, we suggest that if the user’s SNP set of interest is amenable to a reliable statistical ranking, they should consider using tools that make use of this information (69–71).

There are also “topology-based” tools (12) for gene set characterization that take into account the internal structures of pathways, and there is emerging evidence that they may have a practical advantage over methods that ignore pathway topology. Such methods typically make use of additional information about the gene set to be characterized (e.g., expression levels or differential expression statistics), and their extension to SNP set characterization is an interesting but challenging task for future studies. Notably, other authors have investigated the use of network analysis in the context of SNPs but with different goals, e.g., to discover modules or “communities” of inter-related SNPs and genes (72) and perform pathway enrichment tests on these modules (71) or to predict disease-related SNPs (73).

VarSAn allows for special handling of non-coding and coding SNPs in the query set. Since a coding SNP is connected to a gene only if it is predicted to impact the encoded protein’s function (by PolyPhen-2), we made the assumption that multiple coding SNPs linked to the same gene provide proportionate evidence for the gene’s relevance to the biological process represented by the SNP set. On the other hand, multiple non-coding SNPs in the query set connected to the same gene may merit special handling. For instance, if the query set resulted from a GWAS study, they may include multiple non-coding SNPs in the cis-regulatory region of the same gene on account of high linkage disequilibrium (LD), and may not represent proportionate evidence for the gene’s relevance. A similar concern applies when multiple non-coding SNPs are connected to the same gene based on eQTL evidence, since a functional eQTL is likely to be accompanied by eQTLs at segregating positions in LD with it. This is the motivation behind the “SNP-weight” mode of VarSAn, which lets the user down-weight the contribution of multiple non-coding SNPs connected to the same gene. It is a simplistic, optional solution since it does not utilize LD information, a goal left for future work. It does however have the practical advantage that a user does not need to tie their analysis to a specific population group for which LD structure is available.

To some extent, associating a SNP or gene set with a pathway (or a Gene Ontology term) is a subjective goal: the numerous existing methods, especially for gene set characterization, have relied partly on intuitive appeal of the method itself (e.g., the query gene set is enriched for genes of a pathway) (74) and partly on subjective demonstrations of their capabilities (e.g., a cancer-related gene set was most strongly associated with a P53-related pathway) (75). Rigorous evaluation of these methods has been challenging. For instance, Nguyen et al. (12), in one of the most comprehensive efforts to date on benchmarking gene set characterization tools, considered gene sets obtained from expression-based studies of specific diseases and asked how a tool ranks a specific “target pathway” known to be related to the same disease. This approach, while apparently unbiased, does not objectively establish that the assumed “target pathway” is the most desired pathway or how undesirable other pathways are. We believe that the difficulties of benchmarking these tools are tied to the subjective nature of the task itself. In recognition of this fundamental limitation, we resorted to examining complementarity between results from VarSAn and other approaches (HGT, or variants of VarSAn itself), leaving final interpretation to the reader.

At the same time, we did take an important step towards objective benchmarking of different methods for SNP set (or gene set) characterization. Instead of pre-determining the most desirable pathway as a “ground truth”, our evaluation pre-supposes that pathway ranking should be consistent when analyzing different random subsets of the same query set (representing the same disease) and different when analyzing query sets representing different diseases. Our “consistency score” and “confusion score” are heuristics that capture this simple idea, and we assessed a pathway ranking tool by the contrast between these two scores. This avoids commiting to subjective definitions of ground truth. We believe more statistically rigorous implementations of this idea have the potential to move us towards systematic benchmarking for this important bioinformatics task.

Our work on the VarSAn tool has also opened up several opportunities for future research. We mentioned a few of these above, e.g., incorporation of pathway structure, utilization of SNP scores (such as GWAS p-values), and appropriate handling of non-coding SNPs in linkage disequlibrium. Another exciting possibility is to include additional node and edge types in the network in order to integrate new types of relevant information. For example, it may be possible to include gene regulatory information in the form of transcription factor (TF) nodes and edges that connect non-coding SNPs to TFs whose binding sites may be impacted as well as edges from TFs to their potential regulatory target genes. We have also speculated on a more iterative version of propagating information along network edges: once pathways have been scored for relevance to the query set of SNPs, perhaps the SNPs could be scored (also via RWR) for relevance to the top-ranking pathways, and by iterating pathway and SNP ranking multiple times we may identify a subset of the query set that is strongly connected with a subset of the pathways. Finally, in contrast to the “unsupervised” approach adopted in VarSAn, future work may considering deploying supervised learning methods to characterize patterns of network connectivity that are predictive of membership in the query set, and infer pathway relevance scores from these patterns; such an approach was used for gene set characterization in recent work (76).

## AVAILABILITY

Variant Set Annotation (VarSAn) software is available in the Github repository. (https://github.com/UIUCSinhaLab/VarSAn)

## SUPPLEMENTARY DATA

Supplementary Data are available at BioRXiv online.

## ACKNOWLEDGEMENT

We thank Drs. Timothy M. Olson and Jeanne L. Theis at the Mayo Clinic, Rochester, MN for providing whole genome sequencing data from hypoplastic left heart syndrome patient-parent trios under the auspices of the Todd and Karen Wanek Family Program for Hypoplastic Left Heart Syndrome.

