## Supplementary File for "VarSAn: Associating pathways with a set of genomic variants using network analysis"

**Supplementary Materials**: The file contains supplementary figures and tables in the following order.

1. **Supplementary Figure S1:** VarSAn “SNP-weight mode”
2. **Supplementary Table S1 (Separate xls file):** Top pathways reported by VarSAn for breast/prostate cancer (GWAS and somatic mutation) query sets and for HLHS de novo mutation query sets. For results from the two GWAS query sets and for the HLHS query set, literature evidence is provided for most of the top reported pathways. For the four cancer-related query sets, only top 15 pathways are provided (ranked first by empirical p-value and then by equilibrium probability), while for the HLHS query set all significant pathways (empirical p-value < 0.05) are shown.
3. **Supplementary Figure S2:** VarSAn results with SNP-gene network based on prostate tissue eQTL vs Pan-tissue eQTL
4. **Supplementary Figure S3:** VarSAn results compared to alternative where SNP-gene network is based on location
5. **Supplementary Table S2:** Information about disease GWAS SNPs from GWAS Catalog
6. **Supplementary Figure S4:** Consistency and confusion scores of Hypergeometric test (HGT) approach
7. **Supplementary Figure S5:** Direct connections from HLHS de novo variants to VEGFA pathway
8. **Supplementary Figure S6:** Example of indirect connections from HLHS de novo variants to VEGFA pathway
9. **Supplementary Table S3:** Average rank of target pathways at varying noise levels
10. **Supplementary Note S1:** Simplification of REACTOME pathway hierarchy
11. **Supplementary Table S4 (Separate xls file):** REACTOME pathways
12. **Supplementary Table S5 (Separate xls file):** HLHS De Novo mutations
13. **References**

**Supplementary Figure S1**. VarSAn allows for an optional "SNP-weight" mode of execution where each of *N* non-coding SNPs connected to the same gene is assigned a weight of 1/*N* ("NCS*" in panels A,C)*,* while every coding SNP has weight equal to 1 ("CS*" in panels B,C). A SNP node may be connected to multiple genes; in such cases its weight is the sum of its assigned weights due to its connection to each gene (panel D).The RWR algorithm uses these weights during the restart step, hopping back to a SNP node with probability proportional to its weight. Numbers next to gene nodes in all panels indicate the relative importance of that gene in ranking pathways for relevance to the SNP set.

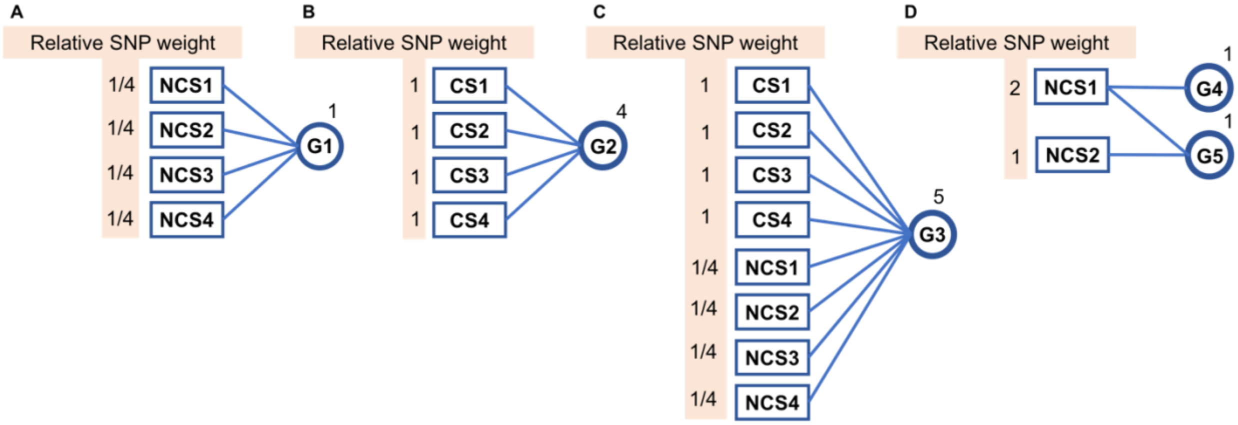

**Supplementary Table S1 (Separate xls file)**. Significant pathways reported by VarSAn for breast cancer GWAS SNPs, breast cancer somatic mutations, prostate cancer GWAS SNPs, prostate cancer somatic mutations and HLHS de novo variants respectively. VarSAn_full was used for query sets comprising GWAS SNPs. For BrCA and PCa somatic mutations that are in coding regions, only “probably damaging” predictions by PolyPhen-2 were used to define SNP-gene edges in the network. For VarSAn analysis of HLHS de novo mutations, SNP-gene edges in the network were based on whether the variant was within 10kb of the gene.

**Supplementary Figure S2**. Scatter plot of pathway rank using VarSAn on the prostate cancer GWAS query set, with SNP-gene edges based on GTEx eQTLs from the relevant tissue (y-axis) versus those based on eQTLs from all tissues (pan-tissue approach, x-axis). The two rankings are correlated with Spearman's rho of 0.29 and p-value of 1.12E-10.

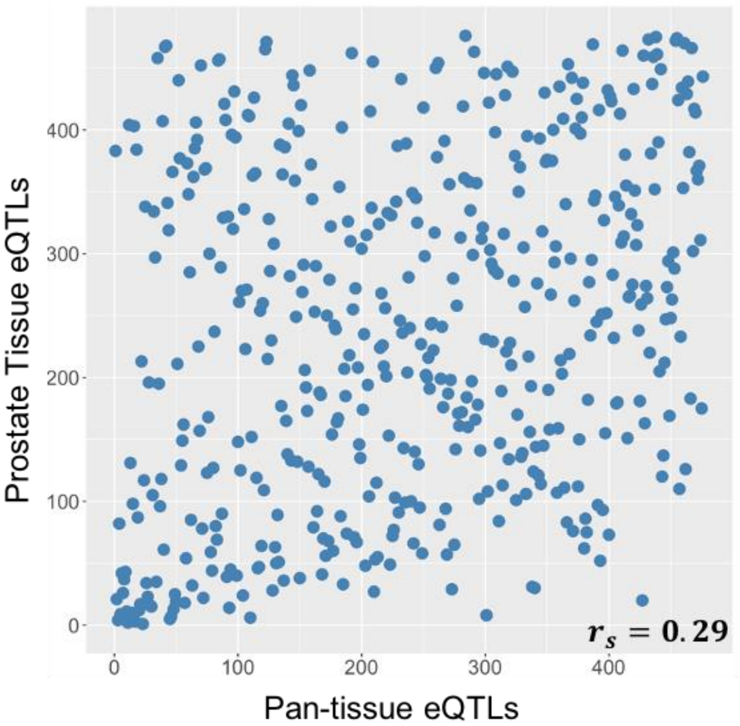

**Supplementary Figure S3**. Using proximity-based SNP-gene connections, VarSAn reports very different pathway ranks (x-axis) than its default settings (y-axis). In this alternative scheme (VarSAn_location), a SNP-gene edge is added to the network if the SNP is within 10k bp of the gene.

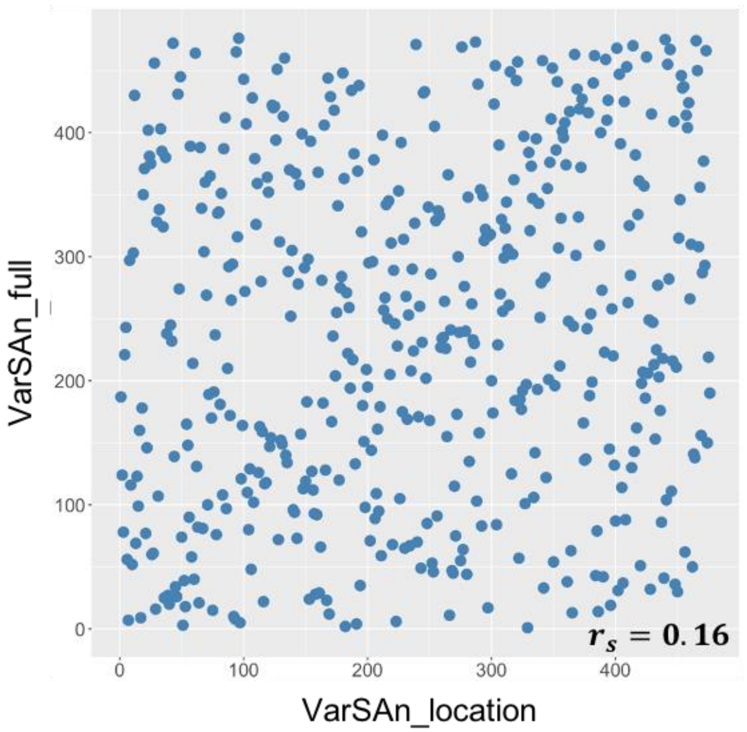

**Supplementary Table S2.** GWAS SNPs for diseases were downloaded from GWAS Catalog (1). GWAS SNPs for diseases were used to evaluate VarSAn's consistency and confusion scores, limiting to diseases for which 200 or more GWAS SNPs were available. (Only GWAS SNPs that could be connected to at least one gene in the SNP-gene network were counted.) The number of GWAS SNPs for each disease, as well as the number of coding and non-coding SNPs are listed in the table below.

| Disease | # GWAS SNPs | # Non-coding SNPs | # Coding SNPs |
|---|---|---|---|
| Asthma | 217 | 214 | 3 |
| Atrial fibrillation | 218 | 216 | 2 |
| Breast cancer | 411 | 399 | 12 |
| Colorectal cancer | 170 | 167 | 3 |
| Crohn disease | 297 | 287 | 10 |
| Inflammatory bowel disease | 270 | 261 | 9 |
| Insomnia | 231 | 229 | 2 |
| Male-pattern baldness | 494 | 486 | 8 |
| Neuroticism | 642 | 638 | 4 |
| Schizophrenia | 739 | 735 | 4 |
| Systemic lupus erythematosus | 339 | 338 | 1 |
| Type 2 diabetes | 511 | 489 | 22 |

**Supplementary Figure S4**. Consistency scores (diagonal values) and confusion scores (off-diagonals) for HGT rankings across all diseases and disease-pairs. The CC ratio, shown at the bottom of the figure, has a median of ~1.9, which is smaller than the median of CC ratio when VarSAn was used for pathway ranking (~3.3).

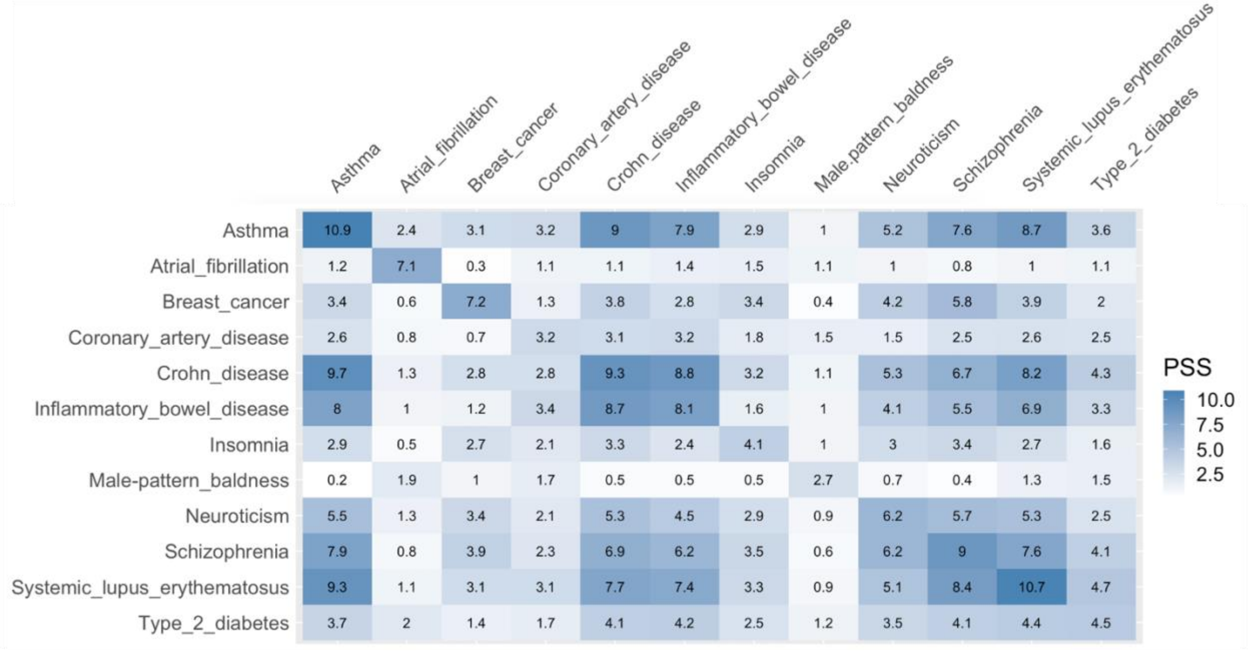


| | | | | | | | | | | | |
|---|---|---|---|---|---|---|---|---|---|---|---|
| **CC-ratio** | 2.20 | 6.17 | 2.87 | 1.48 | 1.90 | 1.90 | 1.71 | 2.84 | 1.72 | 1.96 | 2.21 | 1.55 |

**Supplementary Figure S5.** Direct connections between 10 HLHS de novo variants to VEGFA-VEGFR2 pathway. These variants (left) are located near genes (middle) which are members of the pathway.

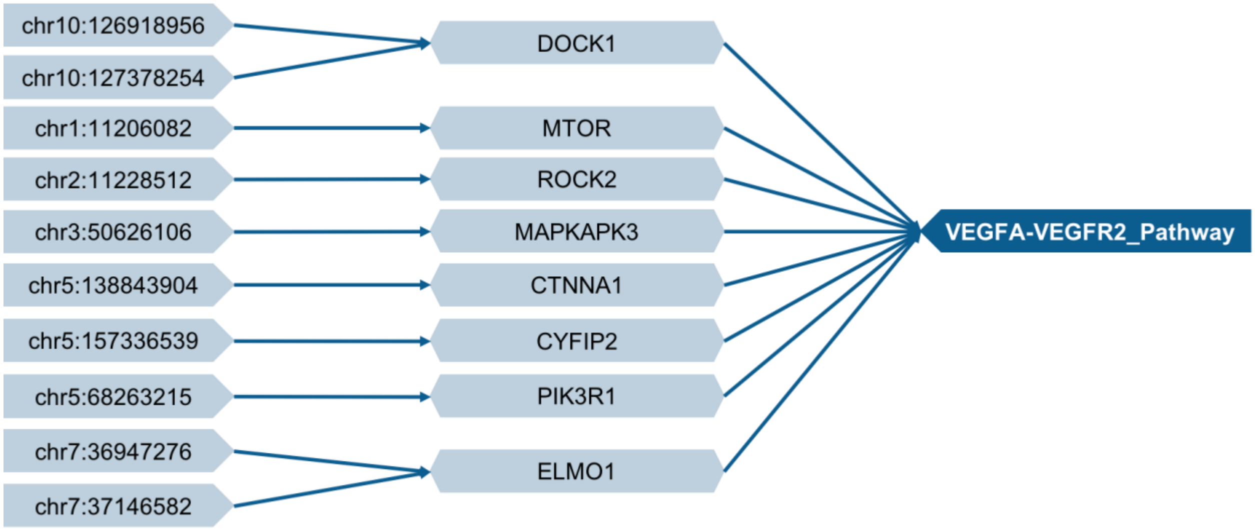

**Supplementary Figure S6.** Examples of HLHS de novo mutations connected to VEGFA-VEGFR2 via indirect connections.

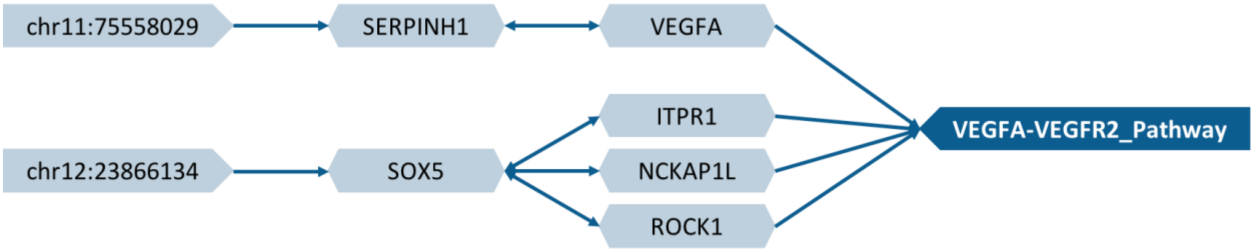

**Supplementary Table S3.** For each drug, shown here is the average rank of its target pathway across 20 independent tests of VarSAn, at four different noise levels. The target pathway for a drug is identified by a VarSAn analysis of the 100 strongest GWAS SNPs for response to that drug. The top five SNPs most connected to the target pathway are then treated as the clean query set, and noisy query sets are created by adding 10x, 20x, 30x or 40x randomly selected SNPs to the query set. VarSAn analysis of the noisy query set is then examined for the rank of the target pathway. This is repeated 20 times for each noise level.

| | **10x** | **20x** | **30x** | **40x** |
|---|---|---|---|---|
| 6MP | 3.2 | 9.05 | 12.65 | 16.05 |
| 6TG | 1.2 | 4.2 | 6.7 | 8.25 |
| ARAC | 1.2 | 2.55 | 6.85 | 11.65 |
| ARSENIC | 3.7 | 5 | 14.05 | 16.95 |
| CARBOPLATIN | 1.2 | 1.45 | 1.4 | 1.85 |
| CDDP | 1 | 1.1 | 1.75 | 1.65 |
| CLADRIBINE | 3.65 | 15.7 | 15.95 | 26.75 |
| DOCETAXEL | 1.25 | 4.05 | 4.65 | 9.95 |
| DOXORUBICIN | 2.95 | 3.95 | 5.7 | 11.2 |
| EPIRUBICIN | 1 | 2.65 | 9.35 | 11.85 |
| EVEROLIMUS | 20.95 | 26.8 | 34.05 | 36.2 |
| FLUDARABINE | 1.5 | 1.7 | 1.65 | 3.8 |
| GEMCITABINE | 6.7 | 17.2 | 23.35 | 39.5 |
| HYPOXIA | 1.2 | 1.4 | 2 | 2.55 |
| METFORMIN | 10.3 | 13.25 | 28.75 | 31.7 |
| MPA | 1.45 | 1.9 | 2.6 | 3.7 |
| MTX | 3.35 | 7.5 | 14.65 | 18.75 |
| NAPQI | 2.9 | 5.1 | 10.35 | 14.85 |
| OXALIPLATIN | 1.2 | 1.4 | 1.15 | 1.55 |
| PACLITAXEL | 2.95 | 9.2 | 11.9 | 12.05 |
| RADIATION | 1.05 | 1.25 | 1.05 | 1.15 |
| RAPAMYCIN | 1 | 4 | 5.95 | 8.55 |
| TCN | 1 | 1 | 1.05 | 1.2 |
| TMZ | 1 | 1.05 | 1.2 | 1.55 |

**Supplementary Note S1.** REACTOME pathways have a hierarchy where the parent pathway is a superset of one or more child pathways, which leads to redundancies among pathway nodes,. To reduce the redundancy between pathways, we only used the pathways at the lowest level of the hierarchy (not a parent for any pathway). For small pathways that have fewer than 30 genes, we replaced them with the first ancestor pathway that has more than 30 genes or the largest ancestor pathway if no ancestory pathway has over 30 genes. Pathways with fewer than 10 genes were excluded.

**Supplementary Table S4 (Separate xls file).** Here we list the name and size of the REACTOME pathways used in this study.

**Supplementary Table S5 (Separate xls file).** Hypoplastic left hear syndrome (HLHS) de novo SNPs from trio-based studies.
